# *‘Candidatus Peptacetobacter felis’* sp. nov., a novel bile acid-converting bacterial species isolated from the feces of healthy cats in the United States

**DOI:** 10.1101/2025.11.28.691207

**Authors:** Niokhor Dione, Kodjovi D. Mlaga, Guillaume Jospin, Zara Marfori, Siyi Liang, Holly H. Ganz

## Abstract

Six strains of a novel anaerobic bacterial taxa exhibiting bile acid-transforming activity were isolated from fecal samples of clinically healthy cats living in Oakland, California. 16S rRNA sequencing and phylogenetic analysis indicate that strain AB800**^T^** belongs to the genus *Peptacetobacter*, with *Peptacetobacter hiranonis* strain JCM 10541 (98% coverage, 99.7% identity) as its closest relative. Whole genome sequencing shows an average nucleotide identity (ANI) of less than 96% (95.73% ANI, 90.53% AF) with the closest validly named species being *Peptacetobacter hiranonis DGF055142*. The size is about 2.58 Mb, containing 2,355 predicted coding sequences and 2,450 annotated genes. Phenotypic characterisation, and comprehensive genomic analysis, including ANI, digital DNA-DNA hybridization (dDDH) and core genome phylogenetic analysis, placing the strain AB800**^T^** on a separate branch, supported the classification of a novel species, strain AB800^T^ (DSM 120482 = LMG 34035) with the proposed name ‘*Candidatus Peptacetobacter felis’* sp. nov., derived from “*felis”* the latin name of cat *felis catus*. These findings expand our understanding of host-associated bile acid converters and provide promising candidates for probiotic development.

## INTRODUCTION

Comprehensive characterisation of the gut microbiome in companion animals and identification of live bacterial taxa are essential for developing next-generation microbial supplements, live biotherapeutic products, and advancing veterinary health [1, 2]. Yet, much of its diversity remains unexplored, especially at the species level. Among the critical microbial processes in the gut, bile acid metabolism plays a central role in regulating host physiology, microbial community structure and interactions, and intestinal homeostasis [3, 4]. In both humans and animals, gut microbes convert primary bile acids into secondary bile acids, modulating lipid absorption, intestinal signalling, and colonisation resistance effects mediated through host receptors such as FXR and TGR5 [5–7]. In dogs and cats, bile acid metabolism has been shown to have profound health implications, with alterations in bile acid profiles linked to gastrointestinal disorders and dysbiosis [5, 8].

To address this knowledge gap, culturomics, a high-throughput, culture-based microbial discovery method, has emerged as a powerful tool [9, 10]. By combining selective anaerobic cultivation, MALDI-TOF MS profiling, and whole genome sequencing, culturomics enables the isolation of novel gut microbes and the functional exploration of their metabolic capabilities [11, 12]. While metagenomics offers broad surveys of genomics content, only culture-based techniques provide direct insight into physiology, metabolic activity, and potential therapeutic applications. The gastrointestinal tracts of healthy cats and dogs provide a rich anaerobic environment that supports diverse microbial consortia involved in key transformations, such as protein fermentation and bile acid conversion [1, 13]. Protein-rich diets common to companion animals enhance microbial fermentation potential, but may also lead to the accumulation of ammonia and other intermediates that disrupt gut stability[14, 15]. In this context, bile acid-transforming bacteria may help stabilise the microbial community and maintain gut homeostasis[5, 16]. One of the main bacteria reported to modulate bile acid levels in companion animals with chronic enteropathy is *Pepetacetobacter (Clostridium) hiranonis* [17].

The *Peptacetobacter* genus is classified in the *Peptostreptococcaceae* family and includes a group of rod-shaped Gram-positive, obligatory anaerobic bacteria. There are two members of the genus *Peptacetobacter*: *Peptacetobacter hiranonis* and *Peptacetobacter hominis*. The first species isolated from this genus was initially classified as *Clostridium* based on the 16s rRNA similarity and DNA-DNA hybridisation analysis [18]. Then, in 2020, the second species was isolated with a ∼95% 16S rRNA gene similarity, and exhibited ANI values of 66–76% (< 96%) and dDDH values of 17–35% (< 70%), far below accepted genus and species thresholds. The whole-genome phylogenomics further demonstrated that these organisms belonged to a distinct lineage, separated from *Clostridium sensu stricto*, leading to the establishment of the genus *Peptacetobacter*, with *P. hominis* as the type species and *P. hiranonis* reclassified accordingly ([19].

Here, we report the isolation and characterisation of a novel bile acid-transforming species from the feces of healthy cats: ‘*Candidatus Peptacetobacter felis’* sp. nov., expanding the genus *Peptacetobacter* to three species. Using a culturomics framework, we identified and analysed these strains through phenotypic, phylogenetic, chemo-taxonomic, and genomic methods. This discovery enhances our ability to conduct additional studies to better understand the bile acid microbiome axis in companion animals. It offers novel candidates for the development of precision probiotics to improve gastrointestinal health and metabolic stability in dogs and cats.

### SAMPLE COLLECTION AND ISOLATION

Fecal samples were collected from five different healthy cats living in Oakland, California, USA, in 2022. Strain AB800**^T^**was isolated from a fecal sample obtained from an 8-year-old, spayed domestic short-haired cat. AB403 was isolated from a fecal sample obtained from a 2-year-old, spayed female domestic short-haired cat. AB804 was isolated from a fecal sample obtained from a 7-year-old, spayed female domestic short-haired cat. AB807 and AB808 were isolated from the same 6-year-old, neutered male American short-haired cat. Strain AB811 was cultivated from a fecal sample obtained from a 1-year-old, spayed female domestic long-haired cat.

Freshly collected samples were transported in BIOME-Preserve AS-930 (Anaerobe Systems, Morgan Hill, CA) transport media in a one-to-one ratio and processed as part of the AnimalBiome culturomics process [20]. Culture and isolation were carried out between September 2022 and February 2023. In an AS500 Anaerobic chamber (Anaerobe Systems, Morgan Hill, CA), the sample was serially diluted, and the diluted samples were plated on Brain Heart Infusion Agar with Horse Blood and Taurocholate (BHIY-HT) (Anaerobe Systems, Morgan Hill, CA) and incubated anaerobically for 48 hours at 37°C. The grown colonies of strain AB800^T^ were repeatedly subcultured on homemade Columbia agar with 5% sheep blood until pure isolates were obtained. Colony identification of the strain was carried out using MALDI Biotyper smart System RUO microflex LT/SH smart with the BIDAL database v12 (Bruker Daltonics, Bremen, Germany) as previously described [20].

### MORPHOLOGICAL, PHENOTYPICAL, AND BIOCHEMICAL CHARACTERISTICS

After isolation, a pure culture of strain AB800^T^ was morphologically characterised by phase-contrast microscopy and Gram staining. Motility was assessed by wet mount, and biochemical profiling was conducted using API ZYM, API 50 CH, and API Rapid ID 32A strips (bioMérieux). The ribosomal protein MALDI-TOF MS spectra profiling was obtained using MALDI Biotyper smart System RUO microflex LT/SH smart with the software flexAnalysis 3.4 (Bruker Daltonics, Bremen, Germany).

On both Chocolate agar (Fig. 1a1) and Brain Heart infusion agar (Fig. 1a2) (Anaerobe Systems, Morgan Hill, CA), strain AB800ᵀ exhibits white-grayish colonies in the form of a star measuring approximately 1 to 2 mm in diameter. The wet mount reveals a non-motile bacterial cell, while the Gram stain shows Gram-positive bacilli with no endospore formation detected (Fig. 1b2). The electron microscopy characterisation shows that strain AB800ᵀ cells measure approximately 0.7–1 µm in width and 3.2–8 µm in length (Fig.1 b1). Growth of AB800ᵀ is optimal on a Chocolate agar (Anaerobe Systems, Morgan Hill, CA) plate at 37 °C in strict anaerobic conditions. The Protein profile characterized by the mass spectra (Fig. 1c) obtained by MALDI-TOF Mass spectrometry, shows a reproducible spectrum among multiple replicates composed of the cytosolic and main ribosomal proteins. This spectrum is unique and composed of peaks within the 2,000 to 15,000 m/z range.

**Fig. 1:**
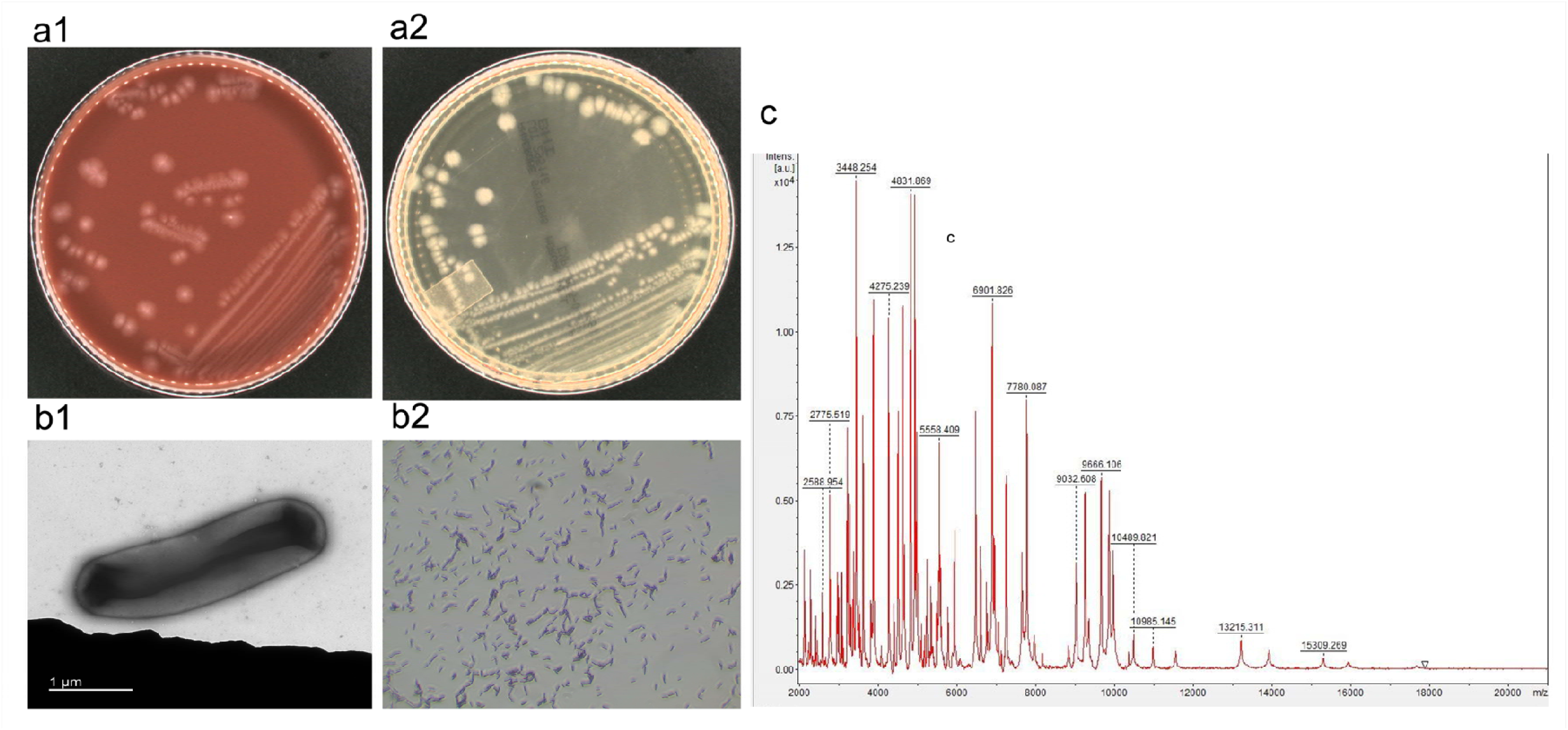
Phenotypic characterization and structure of strain AB800**^T^**. a1) and a2) show colony morphology on Chocolate agar and Brain Heart Infusion agar, respectively (Anaerobe Systems, Morgan Hill, CA); b1) shows transmission electron microscopy structure with a rod-shaped bacterium; b2) represents Gram staining under 100X light microscopy; c) displays the MALDI-TOF spectrum with the main protein profile and associated molecular weight.

The enzymatic activity evaluation shows that strain AB800ᵀ exhibits positive activity for N-acetyl-β-D-glucosaminidase, acid phosphatase, alkaline phosphatase, naphthol-AS-BI-phosphohydrolase, proline aminopeptidase, and esterase lipase (C8). Negative activity was observed for β-glucosidases, β-galactosidase, valine, leucine arylamidases, lipase (C14), α-chymotrypsin, and trypsin. Regarding carbohydrates, strain AB800ᵀ can metabolise D-glucose and D-mannose but shows no activity towards sucrose or sorbitol. Strain AB800ᵀ shows no activity for urea utilisation, nitrate reduction, indole production, and gelatin hydrolysis (Table 1)

**Table 1:** Phenotypic characteristics of strain AB800**^T^**, including morphology, enzymatic, amino acids, salts, utilization and activity, carbon sources of strain AB800**^T^** in comparison with existing members of the genus *Peptacetobacter*, TO-931**^T^**, HD-17, and ZHW00191**^T^**.

| Substrates tested | *Peptacetobacter Felis*<br>AB800<sup>T</sup> | *Peptacetobacter hiranonis*<br>TO-931<sup>T</sup> | *Peptacetobacter hiranonis*<br>HD-17 | *Peptacetobacter hominis*<br>ZHW00191<sup>T</sup> |
| --- | --- | --- | --- | --- |
| **Basic characteristics** |  |  |  |  |
| Cell diameter (µm) |  | 0.8 × 1.6–10 | 0.8 × 1.6–10 | 0.7–1 × 3.2–8 |
| Oxygen requirement | Strict Anaerobe | Strict anaerobe | Obligate anaerobe | Obligate anaerobe |
| Gram stain | + | + | + | + |
| Motility | non-motile | non-motile | non-motile | non-motile |
| Endospore formation | - | + | + | - |
| **Enzyme** |  |  |  |  |
| Alkaline phosphatase | + | NA | NA | + |
| Esterase (C 4) | + | NA | NA | w |
| Esterase Lipase (C 8) | + | NA | NA | + |
| Lipase (C 14) | - | NA | NA | NA |
| Leucine arylamidase | + | NA | NA | NA |
| Valine arylamidase | - | NA | NA | NA |
| Cystine arylamidase | - | NA | NA | NA |
| Trypsin | - | NA | NA | NA |
| α-chymotrypsin | - | NA | NA | NA |
| Acid phosphatase | + | NA | NA | + |
| Naphthol-AS-BI-phosphohydrolase | + | NA | NA | NA |
| α-galactosidase | - | NA | NA | - |
| β-galactosidase | - | NA | NA | NA |
| β-glucuronidase | - | NA | NA | NA |
| α-glucosidase | - | - | - | - |
| β-glucosidase | - | NA | NA | NA |
| N-acetyl-β-glucosaminidase | + | + | + | + |
| α-mannosidase | - | NA | NA | NA |
| α-fucosidase | - | NA | NA | NA |
| **Amino acids / Salts / Carbon source** | - | NA | NA | NA |
| Urea | - | NA | NA | NA |
| L-Tryptophan | - | NA | NA | NA |
| D-Glucose | + | + | + | + |
| D-Mannitol | - | - | - | w (acid) |
| Inositol | - | NA | NA | NA |
| D-Sorbitol | - | - | - | - |
| L-Rhamnose | - | - | - | - |
| D-Saccharose | + | + | + | - |
| D-Melibiose | - | NA | NA | NA |
| Amygdalin | - | NA | NA | NA |
| L-Arabitol | - | NA | NA | NA |
| Temperature range (°C) | 37 | 30–50 (optimum ≈37) | 30–50 (optimum ≈37) | 25–45 (optimum 37–42) |
| pH range | NA | 6.5–9.0 (optimum 7.5–8.0) | 6.5–9.0 (optimum 7.5–8.0) | 5.5–10.0 (optimum 6.5–7.0) |
| NaCl tolerance | NA | Up to ~1% (reported low) | Up to ~1% (reported low) | 0–2% (optimum 0%) |
| Proline aminopeptidase | NA | + | + | + |
| Bile acid 7α-dehydroxylating activity | NA | High activity | High activity | NA |
| Origin | feline gut | Human gut | Human gut | Human gut |

NA = Not available / not tested / data not reported ; w = Weak reaction ; + = Positive reaction / activity present ;
– = Negative reaction/activity absent

### SEQUENCING AND GENOMIC ANALYSIS

The total nucleic acid of strain AB800ᵀ AB403, AB804, AB807, AB808, and AB811 was extracted using the ZymoBIOMICS™ DNA Miniprep Kit for bacterial cells (Zymo Research, Irvine, CA, USA) as previously described [20]. The concentration of the nucleic acid was measured using a Qubit 4 Fluorometer (Thermo Fisher Scientific, Waltham, MA, USA). The strains AB800**^T^**, AB403, AB804, AB807, AB808, and AB811 were sequenced using Illumina technology by SeqCenter (Pittsburgh, PA, United States) as described in their protocol. Illumina sequencing libraries were prepared using the tagmentation-based and PCR-based Illumina DNA Prep kit and custom IDT 10 bp unique dual indices (UDI) with a target insert size of 280 bp. No additional DNA fragmentation or size selection steps were performed. Illumina sequencing was performed on an Illumina NovaSeq X Plus sequencer in one or more multiplexed shared-flow cell runs, producing 2×151 bps paired-end reads. Demultiplexing, quality control, and adapter trimming were performed with bcl-convert1 (v4.2.4).

### PHYLOGENY AND GENOME FEATURES

Sequencing reads were screened for quality control using Fastqc v0.12.1 [21] and filtered with a Phred score >30. We assembled them using SPAdes genome assembler v3.15.5 [22] implemented in shovill (https://github.com/tseemann/shovill) with default parameters. The final scaffolds were screened for quality using CheckM2 v1.0.2 [23] and taxonomically confirmed using GTDB-Tk v2.4.0 (RefDB version r220) [24] and a Blastn search using BLASTN 2.17.0+ [25] against the 16S rRNA database (NCBI). We annotated the genomes using Prokka v1.14.6 [26]. The genome was searched for antimicrobial resistance and virulence genes using Abricate (https://github.com/tseemann/abricate) (version 1.0.0) and the NCBI AMRFinderPlus [27], ARGANNOT [28], ResFinder [28, 29], CARD [30], and MEGARes [31] databases downloaded in November 2025. In addition, we conducted a screening for metabolism pathways associated with bile acid metabolism using gapseq (version 1.3.1) [32] and searched for complete and incomplete pathways.

The strain AB800ᵀ was assembled into 149 contigs, totalling 2.58 Mb in genome size. Compared with the other five strains, AB800ᵀ is in the upper range of genome size, ranging from 2.43 to 2.86 Mb across all strains. The strain AB800ᵀ encodes 2,355 coding sequences and 2,450 total genes, values that fall centrally within the variation seen among other related strains (2,165–2,624 CDS; 2,261–2,719 genes). The strain carries 16 rRNA genes, 78 tRNA genes, and one tmRNA, with three annotated repeat regions, reflecting the typical genomic architecture present across the set. Regarding antimicrobial resistance determinants, strain AB800ᵀ harbors both tetB(P) and tetA(P) genes among the tetracycline group of genes, while no macrolide (ermQ) or lincosamide (lnuP) genes are detected. This resistance gene profile is shared with the strains AB0403, AB804, and AB811. However there were no genes associated to virulence. The genes associated with bile acid metabolism are broadly conserved across all genomes examined, and strain AB800ᵀ encodes the complete set of core *bai* genes, including *baiN_1, baiN_2, baiB, baiE, baiCD_1, baiCD_2, baiF, baiH* and *baiA2*. As in the other strains, *baiN_3, baiN_4* and *baiN_5* are absent (Table 2)

**Table 2:**
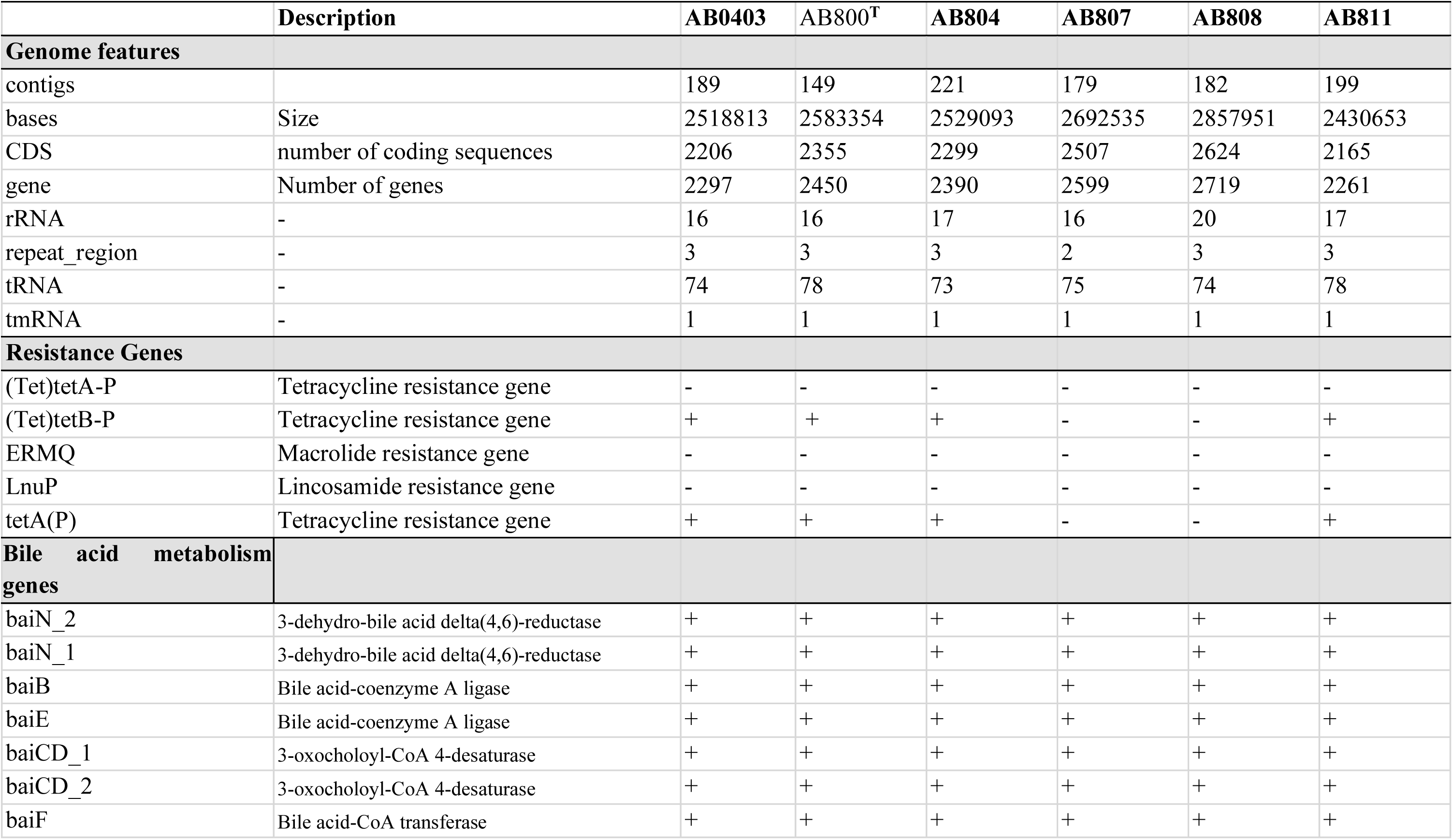
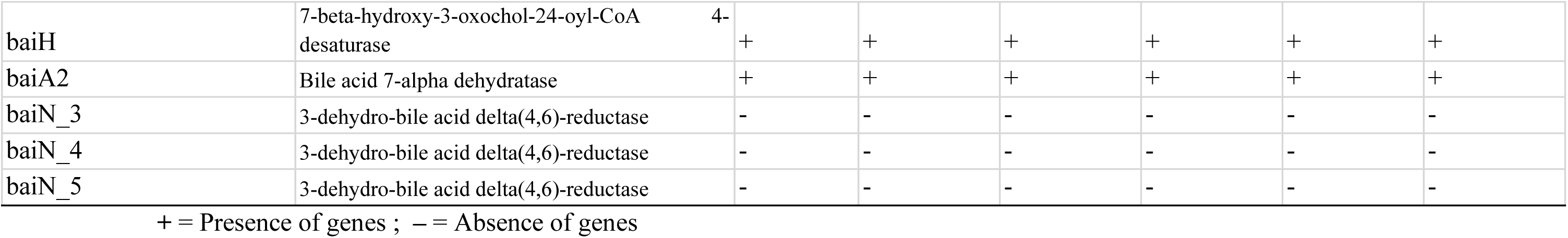
Genomic features of all strains of strain AB403, strain AB800^T^, strain AB804, strain AB807, strain AB808, strain AB811, including antimicrobial resistance genes and genes involved in metabolism pathways.

The 16S rRNA gene sequence of strain AB800**^T^**, when blasted against the 16S ribosomal RNA database, showed 97% coverage and 99% identity to *P. hiranonis JCM 10541*. The maximum likelihood phylogenetic inference with a General Time-Reversible (GTR) model shows strain AB800**^T^** to be a close parent of *P. hiranonis JCM10541(NR178263.1)* and *P. hiranonis strain TO-931 (NR028611.1),* with a bootstrap of 98% and a midpoint rooting topology (Fig. 2: a). Genomic taxonomic classification using Gtdbtk v2 showed that the closest species is *Peptacetobacter* sp900539645 (GCA_900539645.1) with 98.67% ANI, *Peptacetobacter hiranonis (GCF_008151785.1)*, ANI= 94.46%, AF=0.81, and *Peptacetobacter hominis (GCF_006861675.1)*, ANI=80.74, AF=0.45. The DNA-DNA-Hybridization analysis with GGDC showed that the closest from the database is *Peptacetobacter hiranonis DSM 13275* with dDDH = 67.7% (Suppl.Data: Table1). The Core-genome-based phylogeny showed strain AB800**^T^** to be close to or belong to the *Peptacetobacter* cluster but on a separate branch, as *P. hiranonis* and close to the metagenome assembled genomes (MAG) of *Peptacetobacter* (*Pepetacetobacter* sp. bin-44 and *Pepetacetobacter* sp. DOME MAG 11184) (Fig. 2: b). The closest species genomes and other genomes were retrieved from NCBI, and ANI and AF were computed directly using FASTANI (version 1.34) [33]. The results indicated that the closest are *Pepetacetobacter* sp. DOME MAG11184 and *Pepetacetobacter* sp. bin-44 (ANI=98.72%, AF=81.95%, ANI=98.67%, AF=74.17%). The closest known isolate is *Peptacetobacter hiranonis* DGF055145 (ANI=95.73%, AF=81.95%) with ANI < 96%, making strain AB800**^T^** a ‘candidatus’ new species of the genus *Peptacetobacter* (Fig. 3). The 16S rRNA and the genome of strain AB800**^T^** were deposited in Genbank with the respective accession numbers PV358556 and NZ_JBLOUI000000000.1

**Fig. 2:**
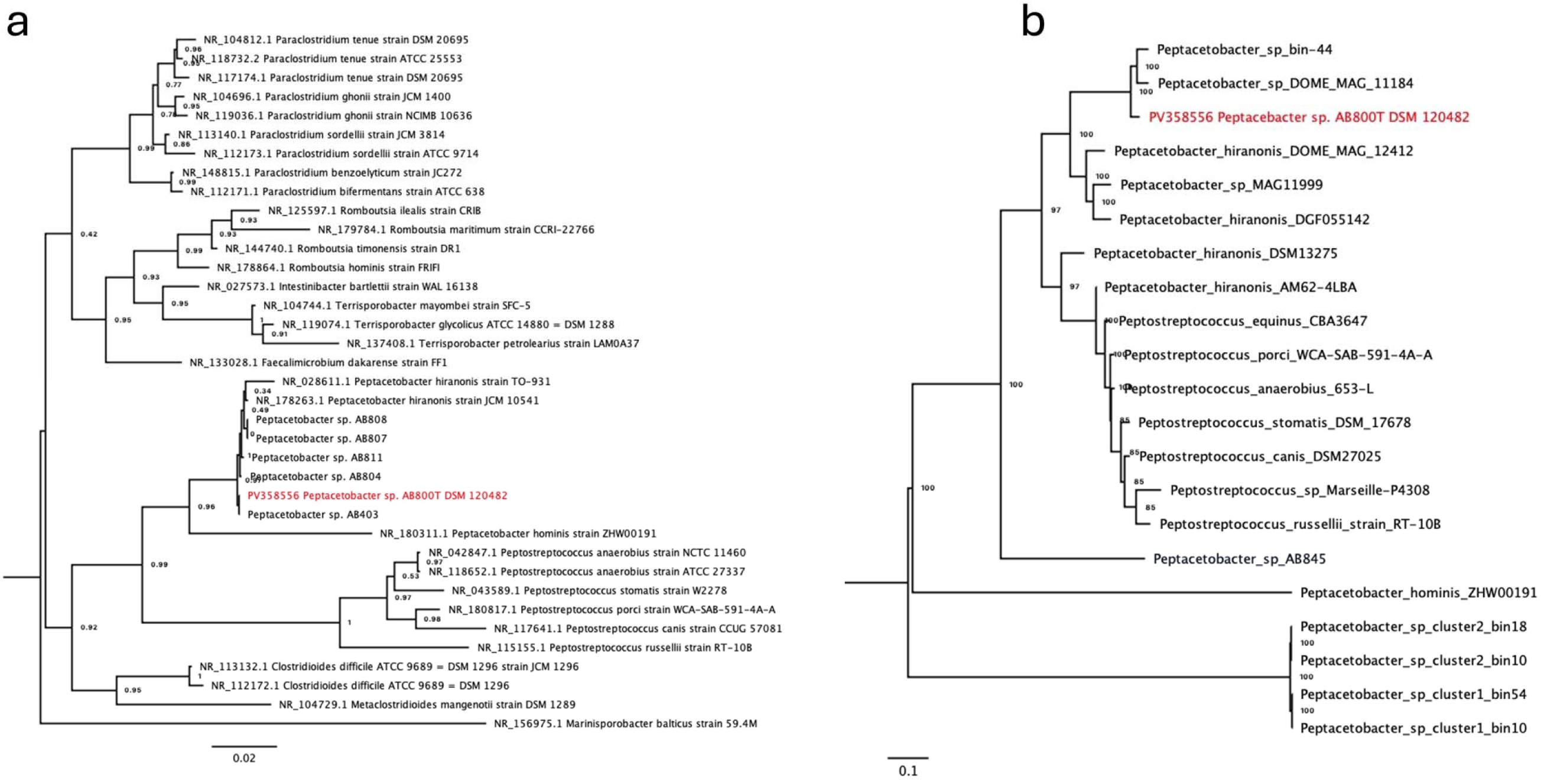
Phylogenetic tree inference using the General Time-Reversible model and bootstrap 1000. **a**: 16S rRNA gene full-length phylogenetic tree with midpoint rooting topology showed strain AB800ᵀ to be close to P. hiranonis. **b**: Core genome alignment phylogeny inference with midpoint rooting, strain AB800ᵀ is shown on a separate branch and close to unknown metagenome assembly species.

**Fig. 3:**
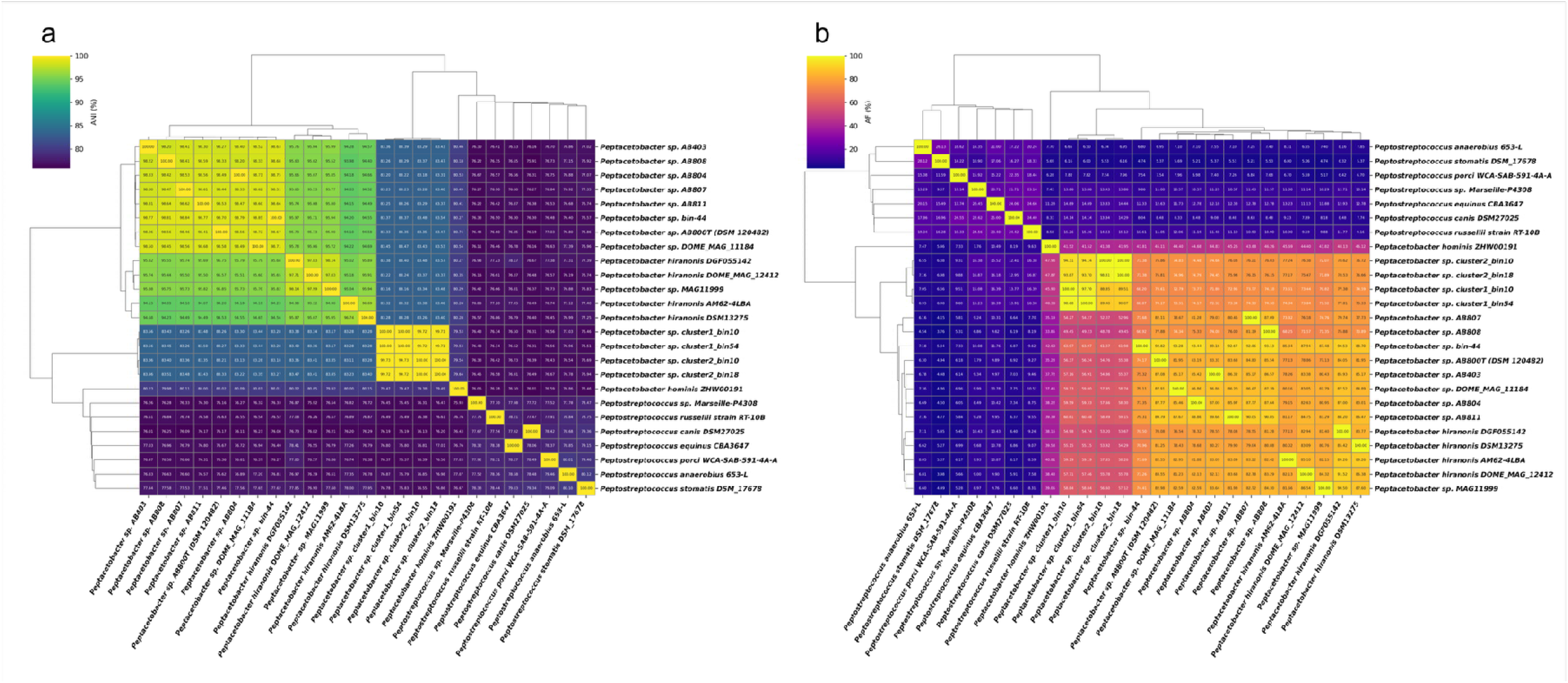
Whole genome taxonomic classification: **a:** Average Nucleotide Identity (ANI) of strain AB800**^T^**compared to closed species retrieved from Gtdbtk classification output. **b:** Alignment Fraction (AF) of strain AB800**^T^** compared to the closest species retrieved from Gtdbtk. The closest species based on the ANI (Average Nucleotide Identity) and Alignment Fraction (AF) are *Pepetacetobacter* sp. DOME MAG11184 and *Pepetacetobacter* sp. bin-44 (ANI=98.72%, AF=81.95%, ANI=98.67%, AF=74.17%). The closest known isolate is *Peptacetobacter hiranonis* DGF055145 (ANI=95.73%, AF=81.95%) with ANI < 96%

### COMPARATIVE GENOMIC

A pangenome analysis using OMCL clustering showed that strain AB800**^T^** shared only 1331 gene clusters with *Peptacetobacter hiranonis* and *Peptacetobacter hominis* (core genome) while conserving 500 genes as unique. It shares 405 with *Peptacetobacter hiranonis* and only 67 with *P. hominis*, confirming that strain AB800**^T^** is closer to *P. hiranonis* than *P. hominis* (Fig. 4: a). The metabolic pathway analysis revealed that, while only *P. hiranonis* and strain AB800**^T^** possess the bile acid 7-alpha and 7-beta dehydroxylation pathways, only strain AB800**^T^** lacks the deconjugation pathways, as shown in Fig. 4: b.

**Fig. 4:**
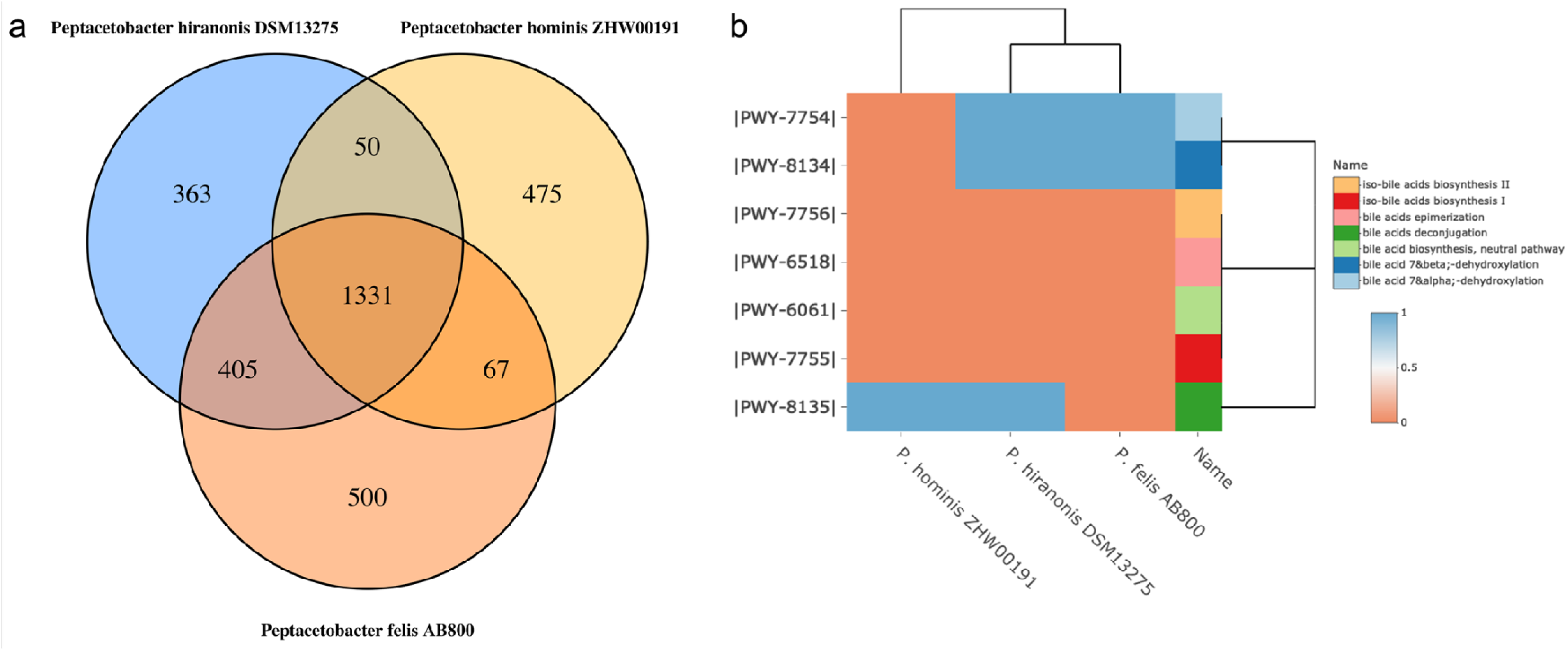
Comparative genomic analysis of strain AB800**^T^** compared to *P. hiranonis* and *P. hominis*. **a**: Venn diagram of the pangenome (ortholog genes) distribution. OMCL clustering showed that strain AB800**^T^** shared only 1331 gene clusters with *P. hiranonis* and *P. hominis,* while conserving 500 genes as unique. It shares 405 with *P. hiranonis* and only 67 with *P. hominis*, confirming that strain AB800**^T^** is closer to *P. hiranonis* than *P. hominis.* **b**: Heatmap showing the distribution of the different components of bile acid metabolic pathways. While only *P. hiranonis* and strain AB800**^T^** possess the bile acid 7-alpha and 7-beta-dehydroxylation pathways, only strain AB800**^T^** lacks the deconjugation pathway

### DESCRIPTION OF *PEPTACETOBACTER SP*. NOV

*‘Candidatus Peptacetobacter felis’* (fe.lis. L. gen. n. felis, referring to the Latin name of cat, felis catus, the species from which it was isolated). A bile acid-converting, Gram-positive, non-motile, obligate anaerobic rod-shaped bacterium isolated from the feces of a healthy cat living in Oakland, California. *Peptacetobacter felis AB800ᵀ* can be grown on Columbia blood agar, Chocolate agar, and Brain Heart infusion agar (with optimal growth on Chocolate agar at 37 °C after 48h. On agar plates, it exhibits white-grayish colonies in the form of a star measuring approximately 1 to 2 mm in diameter. Regarding enzymatic activity, *‘Candidatus Peptacetobacter felis’* AB800ᵀ shows the ability to hydrolyze specific peptide and amino sugar substrates and active phosphatase and esterase systems, which translates to positive results for naphthol-AS-BI-phosphohydrolase, leucine arylamidase, N-acetyl-β-D-glucosaminidase, alkaline phosphatase, esterase (C4), esterase lipase (C8), and acid phosphatase. On the other hand, strain AB800ᵀ does not exhibit activity for cystine arylamidase, trypsin, lipase (C14), valine arylamidase, α-mannosidase, α-fucosidase, α-chymotrypsin, β-glucosidase, or β-galactosidases, showing limited glycosidic activity and proteolytic activity in comparison with other *Peptacetobacter* taxa. Regarding carbohydrate, amino acid, and salt activity, *‘Candidatus Peptacetobacter felis’ AB800ᵀ* is urea- and L-tryptophan-negative, suggesting the absence of urease and tryptophanase activities. For carbon sources, the strain AB800ᵀ can use D-saccharose (sucrose) and D-glucose, but does not use amygdalin, D-melibiose, D-mannitol, D-sorbitol, L-rhamnose, L-arabitol, or inositol. In summary, *‘Candidatus Peptacetobacter felis’ AB800ᵀ* is particularly characterized by its esterase and phosphatase reactivity, but in general by its specific enzyme activities and limited substrate spectrum among the tested compounds. Even though the 16S RNA similarity search and phylogeny did not show a clear delineation between the strain *‘Candidatus Peptacetobacter felis’ AB800ᵀ* and other *P. hiranonis*, Genomic taxonomic classification using Gtdbtk v2 showed that the closest species is Peptacetobacter sp900539645 (GCA_900539645.1) with 98,67% ANI, which is a MAG. However, the nearest known isolate is *Peptacetobacter hiranonis DGF055145* (ANI=95.73%, AF=81.95%), with ANI < 96% and dDDH = 67.7% (< 70%), and Peptacetobacter hiranonis DSM 13275, making *Peptacetobacter sp. AB800T,* a new species of the genus *Peptacetobacter*. The genome is 2,577,873 bp, comprises 149 contigs, has a GC content of 31%, and has been deposited in GenBank under BioProject accession number PRJNA1200056. *‘Candidatus Peptacetobacter felis’* AB800ᵀ was deposited in two international collections with the following accession numbers: *Candidatus Peptacetobacter felis* AB800^T^ (=DSM 120482= LMG 34035

## Ethical statement

The fecal sample used to isolate *‘Candidatus Peptacetobacter felis’* strain AB800ᵀ, as well as the strains AB403, AB804, AB808, AB807, and AB811, were all non-invasively obtained and donated by the cats’ owners with informed consent of the research that will be carried out.

## Author Contributions

ND: Conceived the idea, isolated the strains AB800ᵀ, AB804, AB807, AB808, and AB811, designed the experiments, supervised the characterization experiments, analyzed and validated the data, created the figures, drafted the manuscript, and performed review and preparation. HHG: Conceived the idea, validated the data, reviewed, and prepared the manuscript. KDM: Assembled the genomes, analyzed the data, conducted comparative genomic analyses, drafted the manuscript, and contributed to editing and review. GJ: Prepared the data and deposited the genomes to NCBI, reviewed the manuscript, and contributed to editing. ZM: Isolated strains AB0403, deposited the strains in the international collections (DSMZ and BCCM), and performed phenotypic characterization. SL: Performed the strain purification and cryobanking, DNA extraction, and prepared the DNA for sequencing

## Acknowledgments

We thank the owners of the cats, LizG, Marie, Bwett, Mitz, and Willow, for donating the fecal material used for isolation. We also thank the Animal Biome Microbial Discovery team for assistance with anaerobic culturing and phenotypic characterization, and Seqcenter for sequencing our genomes.

## Supplementary Material

Supplementary Table 1: DNA-DNA Hybridisation table from GGDC

## Data Availability Statement

The genomes of strains AB800^T^, as well as the five strains reported in this study, were submitted and referenced in the NCBI database with the accession numbers JBLOUI000000000 for *‘Candidatus Peptacetobacter felis’* AB800^T^, and the temporary BioSample accession numbers for AB0403, AB804, AB807, AB808, and AB811 are SAMN53260830, SAMN53260831, SAMN53260832, SAMN53260833, SAMN53260834, respectively.

## Abbreviations

MALDI-TOF MS: Matrix-Assisted Laser Desorption/Ionization Time of Flight Mass Spectrometry
WGS: Whole genome sequencing
COG: Clusters of Orthologous Genes
MAG: metagenome assembled genomes
OMCL: Orthologous Markov Clustering

## List of Supplementary data

**Supplementary** Table 1: DNA-DNA Hybridization table from GGDC with strain AB800**^T^** as the

