## Supplementary material for "*‘Candidatus Peptacetobacter felis’* sp. nov., a novel bile acid-converting bacterial species isolated from the feces of healthy cats in the United States": Suplementary table 1

**Supplementary Data**

**Supplementary Table 1**: DNA-DNA Hybridization table from GGDC with strain AB800**^T^** as the query strain

| **Subject strain** | **dDDH**  **(d0, in %)** | **C.I.**  **(d0, in %)** | **dDDH**  **(d4, in %)** | **C.I.**  **(d4, in %)** | **dDDH**  **(d6, in %)** | **C.I.**  **(d6, in %)** | **G+C content**  **difference (in %)** |
| --- | --- | --- | --- | --- | --- | --- | --- |
| *Peptacetobacter hiranonis DSM 13275* | **67.7** | [63.8 - 71.3] | 54.9 | [52.2 - 57.6] | 66.8 | [63.4 - 70.1] | 0.07 |
| *Romboutsia hominis FRIFI* | 13.8 | [11.0 - 17.2] | 23 | [20.7 - 25.4] | 14.1 | [11.7 - 17.0] | 2.64 |
| *Romboutsia ilealis CRIB* | 13.9 | [11.1 - 17.2] | 22.9 | [20.6 - 25.4] | 14.2 | [11.7 - 17.0] | 3.09 |
| *Peptacetobacter hominis ZHW00191T* | 18.5 | [15.4 - 22.1] | 21.9 | [19.6 - 24.3] | 18.2 | [15.5 - 21.2] | 1.74 |
| *Romboutsia weinsteinii CCRI- 19649 T* | 13.2 | [10.5 - 16.5] | 21.5 | [19.2 - 23.9] | 13.5 | [11.2 - 16.3] | 1.65 |
| *Clostridioides difficile DSM 1296* | 13.3 | [10.6 - 16.6] | 21.4 | [19.2 - 23.9] | 13.6 | [11.3 - 16.4] | 2.55 |
| *Intestinibacter bartlettii DSM 16795* | 13.9 | [11.1 - 17.2] | 21.2 | [18.9 - 23.6] | 14.1 | [11.7 - 17.0] | 2.13 |
| *Asaccharospora irregularis DSM 2635* | 13.3 | [10.5 - 16.6] | 21.1 | [18.9 - 23.5] | 13.6 | [11.2 - 16.4] | 0.62 |
| *Paeniclostridium hominis NSJ 45* | 13.7 | [10.9 - 17.1] | 21 | [18.7 - 23.4] | 14 | [11.6 - 16.8] | 3.92 |
| *Terrisporobacter petrolearius LAM0A37* | 13.4 | [10.7 - 16.7] | 20.5 | [18.2 - 22.9] | 13.7 | [11.3 - 16.5] | 2.26 |
| *Romboutsia maritimum CCRI-22766 T* | 13.5 | [10.7 - 16.8] | 20.5 | [18.3 - 22.9] | 13.8 | [11.4 - 16.6] | 3.91 |
| *Eubacterium tenue JCM 6486* | 13.7 | [10.9 - 17.0] | 20.3 | [18.1 - 22.8] | 14 | [11.6 - 16.8] | 4.15 |
| *Intestinibacter bartlettii DSM 16795* | 13.8 | [11.0 - 17.1] | 20 | [17.8 - 22.4] | 14 | [11.6 - 16.9] | 2.32 |
| *Paeniclostridium sordellii ATCC 9714* | 13.4 | [10.6 - 16.7] | 19.9 | [17.7 - 22.3] | 13.7 | [11.3 - 16.5] | 3.56 |
| *Paeniclostridium ghonii DSM 15049* | 13.2 | [10.5 - 16.5] | 19.9 | [17.7 - 22.3] | 13.6 | [11.2 - 16.3] | 3.19 |
